# A mutation in the *LMOD1* actin-binding domain segregating with disease in a large British family with thoracic aortic aneurysms and dissections

**DOI:** 10.1101/153452

**Authors:** Y.B.A. Wan, M.A. Simpson, J.A. Aragon-Martin, D.P.S. Osborn, E. Regalado, D.C. Guo, C. Boileau, G. Jondeau, L. Benarroch, Y. Isekame, J. Bharj, J. Sneddon, E. Fisher, J. Dean, M.T. Tome Esteban, A. Saggar, D. Milewicz, M. Jahangiri, E. R. Behr, A. Smith, A. H. Child

## Abstract

We describe a mutation in *LMOD1*, which predisposes individuals to thoracic aortic aneurysms and dissections in a large multi-generation British family. Exome variant profiles for the proband and two distantly related affected relatives were generated and a rare protein-altering, heterozygous variant was identified, present in all the exome-sequenced affected individuals. The allele c.1784T>C, p.(V595A) in *LMOD1* is located in a known actin-binding WH2 domain and is carried by all living affected individuals in the family. *LMOD1* was further assessed in a consecutive series of 98 UK TAAD patients and one further mutation was found, yielding an incidence of ∼2% in our study group. Assessment of *LMOD1* in international TAAD cohorts discovered nine other missense variants of which three were classed as likely pathogenic.

Validation of *LMOD1* was undertaken using a zebrafish animal model. Knock-down of both *lmod1a* and *lmod1b* paralogs using morpholino oligonucleotides showed a reproducible abnormal phenotype involving the aortic arches under off-target controls. Injection of the human *LMOD1* c.1784T>C, p.(V595A) mutation demonstrated a likely dominant negative effect and illustrated a loss of function cause.

Mutations found in the WH2 actin-binding domain of *LMOD1* may delay actin polymerization and therefore compromise actin length, dynamics and interaction with myosin in the smooth muscle contraction pathway.

## Introduction

TAAD (thoracic aortic aneurysm and dissection) is a severe disease accounting for approximately 15,000 deaths per year in the USA^1^. TAAD can be present in genetic syndromes such as Marfan and Loeys-Dietz syndromes and the vascular form of Ehlers Danlos syndrome. However, most TAAD patients do not satisfy the characteristic syndromes^2^. Approximately a quarter of non-syndromic forms of TAAD are familial^3^. Within affected kindred the disorder shows variability in penetrance and severity, notably between males and females^3–5^. The underlying molecular genetics of this disorder is heterogeneous, with 36 genes reported to date as sites of pathogenic mutation in syndromic and non-syndromic TAAD (OMIM). Mutations in reported genes contribute to the genetic aetiology of approximately 20-30% of TAAD families^6^, with the most mutations, in approximately 14% of families, found in *ACTA2*. Investigations into the biological roles of this group of 36 genes^7–37^ have revealed some common pathways, including those relating to TGF-β signaling^38^ as well as smooth muscle contraction^7^.

To extend the understanding of the genetic basis of TAAD, a whole exome sequencing approach was used in a large multigenerational family of British ancestry (M441) with autosomal dominant inheritance of TAAD. The proband initially consulted a cardiologist as he had lost both his father and half-brother to type A dissection involving the ascending aorta. Doppler transthoracic echocardiography (TTE) measurements revealed he had a mildly dilated aortic root (4.1cm). Clinical investigation of other family members revealed five living relatives who had previously undergone surgery for TAAD or who had a dilated aortic root (figure 1). Physical examination of these subjects revealed that they shared some phenotypic features. Family history revealed aortic dissection as the cause of death for four males (mean age of death 52.25 years, range 39-62 years).

**Figure 1.**
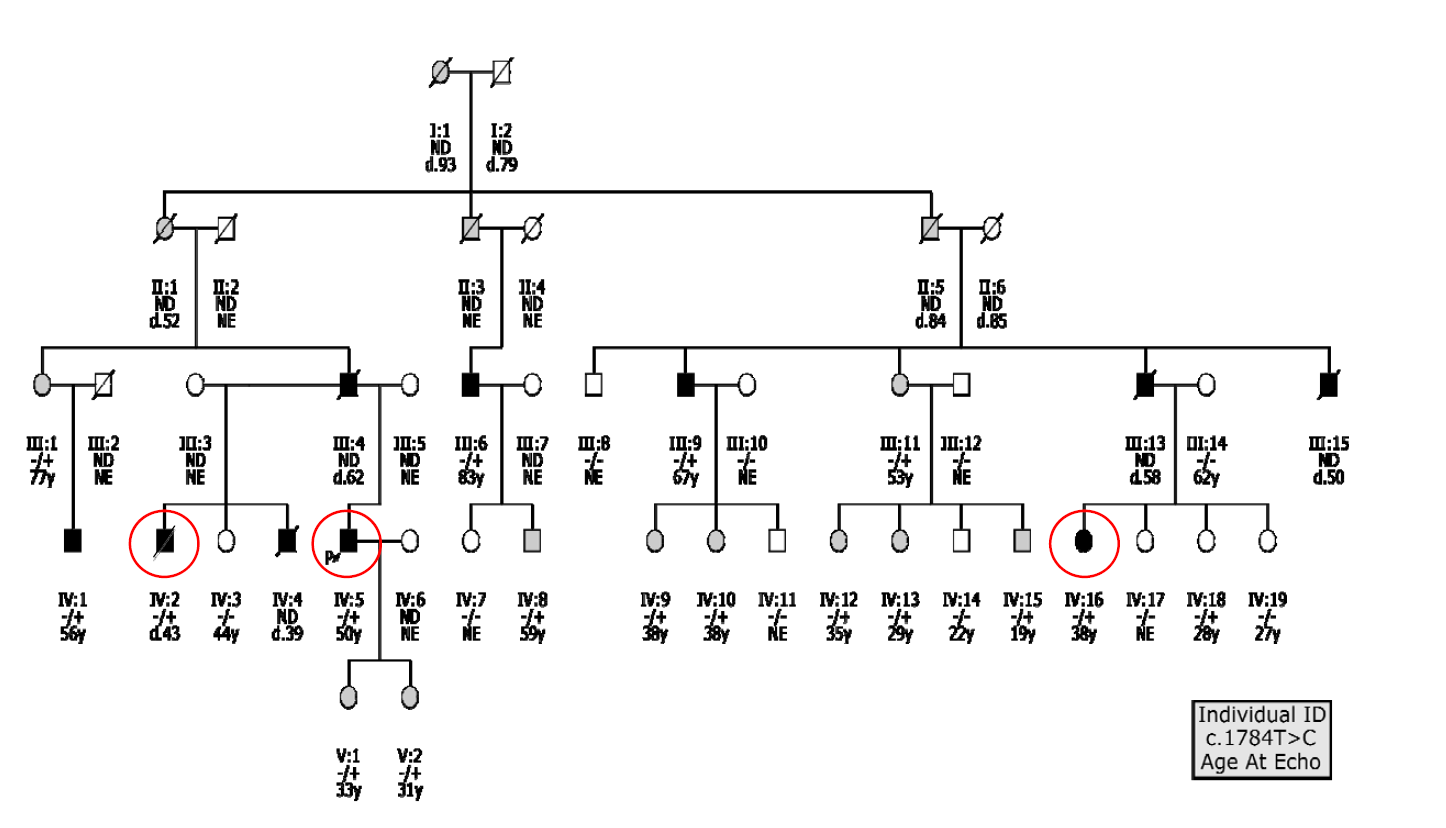
Family M441 pedigree showing affected individuals in black, ambiguous phenotype in grey and unaffected in white. The proband is indicated with a P with an arrow. Affected individuals, who have undergone exome sequencing, circled in red. Individuals carrying c.1784T>C allele (-/+); reference allele (-/-). ND-No DNA; NE-No echo; --y (e.g. 20y)-age when echo was taken (e.g. 20 years old); d.-- (e.g. d.39) age at death (e.g. death at age 39).

## Subjects and Methods

### Subjects

All individuals in this study were ascertained through the Aortopathy clinic at St George’s University Hospitals Foundation NHS Trust.

Ethical consent has been obtained from the National Research Ethics Service Committee London-Bloomsbury (reference; 14/LO/1770, project ID; 99752). Letter of invitation to participants, participant information sheets, DNA analysis consent forms and skin biopsy consent forms have all been approved by the ethical committee. License has been granted for the storage of samples under section 16 (2) (e) (ii) of the Human Tissue Act 2004 (No. 12335). Patient and family participants have read and signed the relevant consent forms.

### Sequencing

We undertook a whole exome sequencing (WES) approach to identify the causative mutation in this family, generating exome variant profiles for the proband and two distantly affected relatives (figure 1). It was performed at the KCL-GSTT BRC Genomics facility at Guys Hospital, London as a collaborative research service. Exome libraries were generated using Agilent SureSelect kits on an Agilent Bravo Liquid handling robot and the exome enriched libraries were sequenced by paired-end reads on an Illumina HiSeq2000 platform and interpreted using their own in-house pipeline.

### Variant Interpretation

In-silico prediction tools Polyphen-2^39^, GVGD^40^, UMD Predictor^41^, SIFT^42^, and PROVEAN^43^ were used along with the genomic database The Genome Aggregation Database (gnomAD)^44^ to determine pathogenicity of alleles found upon exome analysis. Functional annotation tools Combined Annotation-Dependent Depletion (CADD)^45^ and PhastCons 100 way^46^ were used to predict if variants were amongst the top 1% of deleterious variants and estimate the probablility that the nucleotide belongs to a conserved element based on multiple alignment. The Universal Protein Resource (UniProt)^47^ and The Genotype Tissue Expression (GTex)^48^ were used to assess domain architecture and tissue specific gene expression. Sanger sequencing was performed to determine cosegregation of candidate genes in the study family as well as detect further mutations in our consecutive 98 TAAD cohort.

### TAAD gene analysis

Exome profiles were screened for disease-causing mutations in all 36 TAAD associated genes. These genes included *ACTA2* (OMIM: *102620), *BGN* (OMIM: *301870), *COL1A1* (OMIM: *120150), *COL1A2* (OMIM: *120160), *COL3A1* (OMIM: *120180), *COL4A5* (OMIM: *303630), *COL5A1* (OMIM:*120215), *COL5A2* (OMIM: *120190), *EFEMP2* (OMIM: *604633), *ELN* (OMIM: *130160), *FBN1* (OMIM: *134797), *FBN2* (OMIM: *612570), *FLNA* (OMIM: *300017), *FOXE3* (OMIM: *601094), *SLC2A10* (OMIM: *606145), *JAG1* (OMIM: *601920), *LRP1* (OMIM: *107770), *MAT2A* (OMIM: *601468), *MFAP5* (OMIM: *601103), *MYH11* (OMIM: *160745), *MYLK* (OMIM:*600922), *NOTCH1* (OMIM: *190198), *PKD1* (OMIM: *601313), *PKD2* (OMIM:*173910), *PLOD1* (OMIM: *153454), *PRKG1* (OMIM: *176894), *SKI* (OMIM:*164780), *SMAD2* (OMIM: *601366), *SMAD3* (OMIM: *603109), *SMAD4* (OMIM: *600993), *TGFB2* (OMIM: *190220), TGFB3 (OMIM: *190230), *TGFBR1* (OMIM: *190181), *TGFBR2* (OMIM: *190182), *TGFBR3* (OMIM:*600742) and ULK4 (OMIM: *617010).

### Animal studies

Ethics Statement Animal maintenance, husbandry, and procedures are defined and controlled by the Animals (Scientific Procedures) Act 1986. All animal experimentation has been carried out under licenses granted by the Home Secretary (PIL No. 70/10999) in compliance with Biological Services Management Group and the Biological Services Ethical Committee, UCL, London, UK.

Wild type (AB x Tup LF) and Tg(kdrl:GFP) zebrafish were maintained and staged as previously described by Westerfield et al^49^. Tg(kdrl:GFP) fish were used to study the development of the aortic arches^50^.

Zebrafish (*Danio rerio*) were used to assess the involvement of *LMOD1* as a potential TAAD-causing gene as this species have proven to be a versatile vertebrate model to study human genetic disease and have already been used to demonstrate pathogenicity of human TAAD genes^20; 34^. *LMOD1* in zebrafish identifies two paralogs, *lmod1a* and *lmod1b*. Gene duplication is a common feature in zebrafish and these paralogs are the result of a genome duplication that occurred in teleost fish some 300 million years ago^51^. Whole mount in-situ hybridization was performed to determine expression of candidate genes. Morpholino oligonucleotides (MO) for splice blocking *lmod1a* exon1/intron1 (5’-TATCATCATCATCACCTCTGCGCTC-3’) and intron1/exon2 (5’-CCTCCCTACAAACAGACACATTATT-3’) and splice blocking *lmod1b* (5’-ATTTCCATTTCCTACCTTTTTGAGC-3’) were injected into 1-2 cell embryos to knock down candidate genes and evaluate the phenotype of morphants compared to un-injected wild-type control fish using confocal microscopy at 3-4 days post fertilization (dpf). Fluorescent microscopy was employed to investigate MO injected fish processed with an actin-filament phalloidin stain and Dextran dyes, highlighting the vasculature. To obtain an enhanced resolution at confocal microscopy, *Tg(kdrl:GFP)^50^* fish were used to highlight the vasculature, in particular the aortic arches of MO injected fish. A standard control MO (5’-CCTCTTACCTCAGTTACAATTTATA-3’) targeting a splice site of human beta-globin^52^ was used to identify toxicity concentrations. A zebrafish control MO targeting *p53* (5’-GCGCCATTGCTTTGCAAGAATTG-3’) was injected singularly (6ng/ul) as well as co-injected with *lmod1a*/*lmod1b* MOs (6ng/ul total, 2ng/ul + 2ng/ul +2ng/ul) as MOs have been reported to cause off-target activation of p53 and therefore causing non-specific phenotype in injected fish^53^.

Injection of full length human *LMOD1* mRNA and zebrafish *lmod1a* and *lmod1b* at 1-cell stage were performed to rescue the phenotype caused by MO knockdown. Injection of the c.1784T>C mRNA from the study family mutation at 1-cell stage was performed to assess the phenotype of the embryos.

## Results

Initial analysis of exome variant call format (VCF) files for all 36 reported TAAD genes showed no pathogenic mutation present in all three affected individuals. Further analysis revealed three rare (MAF < 0.0001), protein-altering, heterozygous variants present in the profiles of each of the sequenced individuals. The three candidate variants located in *LMOD1*, *SPHKAP* and *ZNF483* were each evaluated for cosegregation with the trait in the pedigree by Sanger sequencing. Neither the variant in *SPHKAP* nor the variant in *ZNF483* cosegregated with the disease. However, the c.1784T>C, p.(V595A) (NM_012134) allele in *LMOD1* was carried by each of the five affected living individuals (figure 1) and was also present in the DNA sample obtained from a deceased affected half-brother of the proband. A further 10 heterozygous individuals were identified in the pedigree who had an ambiguous phenotype, meaning that none of them had detectable aortic dilation by transthoracic echo but they presented with two or more minor features associated with a marfanoid phenotype^54^. Seven of these 10 individuals shared phenotypic features found in the family members who had previously undergone surgery for TAAD or who had a dilated aortic root. One heterozygous clinically unaffected individual was found who did not present with a detectable aneurysm, but this individual, IV:18, aged 28 at the time of examination, was documented as having only a high arched palate consistent with the phenotypic presentation in the proband and 10 other heterozygous individuals. A parametric linkage analysis of the observed segregation of the *LMOD1* c.1784T>C allele and the affection status, generated a LOD score in excess of 2.3, p=0.0005. (LOD score 6.7 from an affected-only analysis,dominant model, disease frequency 0.0001, penetrance 0.9). These data suggested that the allele located in *LMOD1* was likely to be disease-causing in this family and should therefore be investigated further.

The c.1784T>C *LMOD1* variant is located in the third and terminal exon of the encoded 1839bp transcript (figure 2). The predicted p.(V595A) substitution in the mature protein product replaces a valine residue that has been highly conserved throughout evolution. This residue is located in the C-terminal WH2 domain known to be an actin-binding motif. The c.1784T>C allele was not observed gnomAD and was not seen in ˜5,500 exomes sequenced and processed using the same laboratory protocols and data processing pipeline as used in the current study. The rarity of this allele suggests that it must have a role in the pathogenesis of aortic aneurysm in a population of TAAD patients.

**Figure 2.**
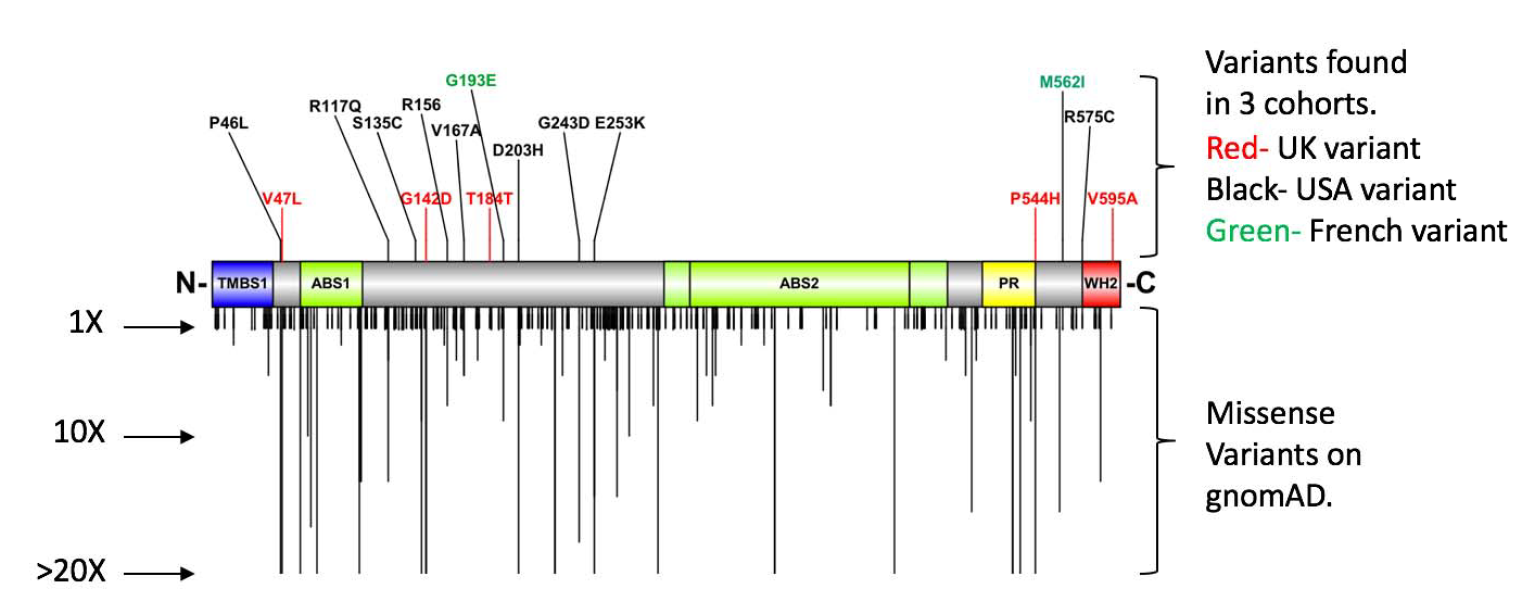
Diagram representing domain architecture of *LMOD1*. TMBS1-Tropomyosin binding site 1 (blue area). ABS1 and ABS2-Actin binding site 1 and 2 (green areas). PR-Proline rich region (yellow area) and WH2-Wiskott-Aldrich syndrome homology region 2 (red area). Adapted from Boczkowska et al, 201563. Upward lines indicate alleles found in three cohorts (red variants from the UK cohort, black from the USA and green from French cohort).Downward lines indicate variants present on gnomAD, and the length indicates the number of times it has been reported from their databa e. Downward lines truncated from 20X.

The possibility that further rare *LMOD1* alleles may explain additional cases of TAAD was explored using Sanger sequencing of the three *LMOD1* exons and their associated splice sites in a cohort of 98 (36/98 familial) unrelated British individuals with TAAD. Sanger sequencing of the three amplicons identified three of the 98 individuals carrying heterozygous missense alleles in *LMOD1* and one heterozygous silent allele (table 1 showing only rare missense alleles and figure 2 showing all variants found). The observation of three missense alleles in this gene in a total of 196 alleles does not deviate from the expected rate based on allele counts in the ExAC (Chi Square p = 0.49). However, it is noteworthy that one proband, individual M508.1, carries a heterozygous c.1631C>A, p.(P544H) (MAF = 0.0004342, rs202184893) allele, which was also predicted to be damaging by all *in-silico* prediction tools used (table 1). This residue substitution is highly conserved across species and appears within the proline-rich region found upstream of the WH2 domain of the protein. This affected female (M508.1) at age 32 had a type A dissection at 36 weeks of pregnancy necessitating emergency caesarean section and aortic root and valve replacement. Aortic histology revealed cystic medial necrosis found typically but not specifically in Marfan syndrome patients. Physical examination revealed individual M508.1 to have a high palate, overcrowded teeth, wrinkled forehead, and small joint hypermobility similar to features seen in the proband and affected members of our initial proband’s family.

**Table 1.** *In-silico* analysis of only the rare missense alleles detected in 98 UK, 800 USA and 403 French TAAD probands. The table includes columns for the cohort, mutation at nucleotide level, consequence on protein level (HGVS-Human Genome Variation Society nomenclature-NM_012134), Combined Annotation-Dependent Depletion (CADD) score, PhastCons score, *In-silico* predictions (Polyphen-2, GVGD, UMD-Predictor, SIFT and PROVEAN (+) - Damaging in two or more in-silico tools used (-) - Damaging in one or none of the in-silico tools used.), gnomAD allele frequency, cosegregation tested (N/A-No cosegregation performed). Rows highlighted in grey are predicted to be disease-causing mutations due to high CAAD scores (>20), PhastCons 100 score (>0.9) and damaging (+) upon in-silico predictions.

| Cohort | Mutation<br>HGVS | Consequence<br>HGVS | CADD<br>score | phastCons<br>100 score | *In-silico*<br>predictions | gnomAD<br>allele<br>frequency | Cosegregation<br>tested |
| --- | --- | --- | --- | --- | --- | --- | --- |
| UK | c.1631C>A | p.(P544H) | 25.8 | 1 | + | 0.0003 | Yes |
| UK | c.1784T>C | p.(V595A) | 23.6 | 0.913 | + | 0 | Yes |
| USA | c.137C>T | p.(P46L) | 23.4 | 1 | + | 0.0003 | N/A |
| USA | c.350G>A | p.(R117Q) | 18.26 | 0 | - | 0.00002 | N/A |
| USA | c.404C>G | p.(S135C) | 24.4 | 0.003 | + | 0.00001 | N/A |
| USA | c.466C>T | p.(R156W) | 25.9 | 0.01 | + | 0.00005 | N/A |
| USA | c.500T>C | p.(V167A) | 11.17 | 0.996 | + | 0 | N/A |
| USA | c.607G>C | p.(D203H) | 23.6 | 0.008 | + | 0.0001 | N/A |
| USA | c.728G>A | p.(G243D) | 0.102 | 0 | + | 0.0002 | N/A |
| USA | c.757G>A | p.(E253K) | 8.519 | 0 | - | 0.0001 | N/A |
| USA | c.1723C>T | p.(R575C) | 33 | 0.999 | + | 0 | N/A |
| FRANCE | c.578G>A | p.(G193E) | 0.016 | 0 | + | 0.0002 | N/A |
| FRANCE | c.1686G>A | p.(M562I) | 23.7 | 1 | + | 0.000008 | N/A |

Individual M508.1, a proband from the consecutive TAAD series reported her maternal grandmother died at age 40 of ruptured dissecting aortic aneurysm, verified by the death certificate. DNA was obtained from the proband’s mother and Sanger sequencing confirmed the c.1631C>A allele was present, although at age 55, after two pregnancies, her echocardiogram was reported to be within normal range. In this family, the proband’s mother appears to be unaffected by aortic disease at present based on echocardiogram measurements, however is physically inactive due to an unrelated illness causing an inflammatory neurological condition. This may be a contributing factor to the variable expressivity seen in this family. The proband’s clinically unaffected sister, whose cardiac evaluation including echocardiogram is normal, does not carry the mutation. Clinical evaluation suggests similarity in phenotypic presentation of affected individuals in both UK families.

The phenotypic presentation seen in affected individuals in both families M441 and M508 are generally ’soft signs’ also seen in normal control populations. Arm span greater than height, high arched palate, blue sclerae, down-slanting palpebral fissures, wrinkled forehead, small joint hypermobility and hyperextensible skin were recorded and compared with the same findings in all our TAAD probands. Only the presence of small joint hypermobility was found to be significantly higher in family M441 and proband M508.1 compared to our TAAD proband cohort (p=0.0000003).

Comparison of our TAAD study family with our cohort of 98 consecutive TAAD probands (table 2), reveals that only blue sclerae and small joint hypermobility appear significantly more often in the M441 family. These clinical features were not assessed in international cohorts as they were not recorded upon physical examination.

**Table 2.** Phenotype correlation between group A-c.1784T>C affected status (n=18) with group B-all clinically diagnosed TAAD probands (n=98). AS>HT-Arm span greater than height, HP-High palate, BS-Blue sclera, RE-Redundant eyelids, DSPF-Downward-slanting palpebral fissures, WF-Wrinkled forehead, SJH-Small joint hypermobility and HS-Hyper extensible skin.

| Study group | AS>HT | HP | BS | RE | DSPF | WF | SJH | HS |
| --- | --- | --- | --- | --- | --- | --- | --- | --- |
| Group A (Family M441 c.1784T>C carrier >18yrs) | 12/18 | 12/18 | 6/18 | 2/18 | 3/18 | 6/18 | 9/18 | 9/18 |
| Group B (All TAAD probands) | 51/98 | 48/98 | 3/98 | 11/98 | 11/98 | 18/98 | 24/98 | 45/98 |
| Chi Square between group A (c.1784T>C carrier >18years) and group B (all TAAD probands) P= | 0.25 | 0.17 | 0.00001 | 0.43 | 0.51 | 0.15 | 0.0000003 | 0.7 |

The incidence of mutations in *LMOD1* was also evaluated in international familial TAAD cohorts; GenTac Research Consortium (800 TAAD families) and Paris Marfan Syndrome Study Group (225 families and 178 isolated cases), which revealed further variants in this gene (Table 1). Nine rare missense variants were detected in 12 familial probands of the GenTac Research Consortium and two rare missense variants were identified in the Paris Marfan Study Group. All heterozygous alleles were found spanning across all three exons of the gene. Variants from all three cohorts were evaluated as likely pathogenic if they were scored in the top 1% of most deleterious substitutions, high conservation and damaging in 2 or more in-silico tools (table 1, highlighted in grey).

Further investigations through the use of *in-vivo* models were performed to show that a non-functional *LMOD1* is associated with TAAD in the zebrafish animal model. Conservation of the human LMOD1 (P29536) WH2 domain at residues 574-600 results in 62.5% shared identity to the predicted zebrafish *lmod1a* (F1Q5S5) and 84.6% to zebrafish *lmod1b* (E7F062) and conservation of the proline-rich region at residues 509-544 results in 51.2% and 56.8% shared identity with *lmod1a* and *lmod1b*, respectively.

These data suggest that zebrafish *lmod1b* is more closely related to human LMOD1 and thus offers the better gene to study, however both paralogs are likely to be expressed and have overlapping functions, therefore functional redundancy is likely to exist.

*In-situ* hybridization revealed strong expression of both *lmod1a* and *lmod1b* in the heart, fin buds and as well as expression from the telencephalon to the tegmentum of the zebrafish. To determine the functional involvement of these genes during development, antisense Morpholino oligonucleotides (MO) were employed that block gene-specific translation or mRNA processing. Gene knockdown was assessed by co-injection of two non-overlapping splice-directed MOs in *lmod1a*, and one splice-directed MO in *lmod1b*. The zebrafish embryos were classified as normal if the phenotype was indistinguishable from wild-type un-injected embryos, moderately affected if they had pericardial oedema and tail curvature, and severely affected if widespread oedema was seen with severe tail curvature (figure 3A). This classification was confirmed independently by two trained observers. Additional phenotypic features observed in the injected morphants included smaller eyes and body size, abnormal distribution of melanocytes compared to wild type un-injected controls (figure 3A).

**Figure 3.**
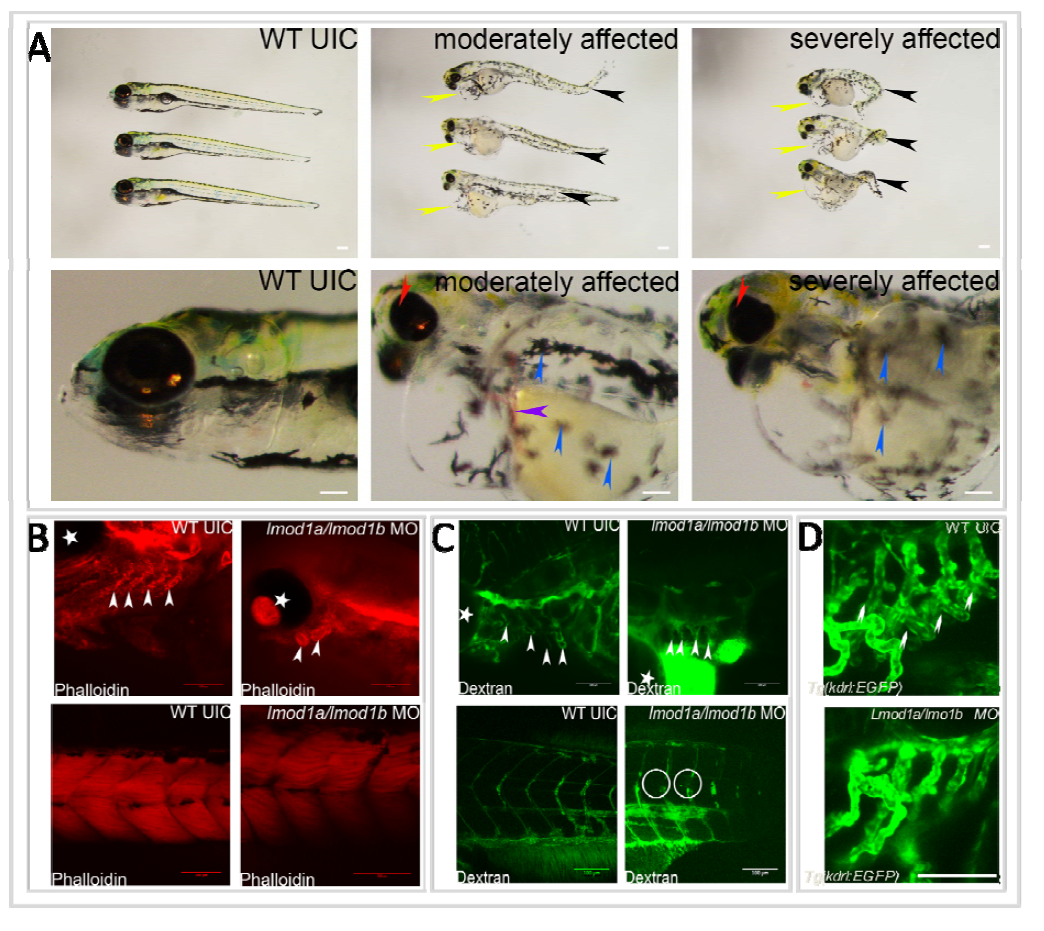
Microscopy of control and *lmod1a*/*lmo1b* MO fish. **A-** Phenotypic representation of un-injected wild-type zebrafish compared to moderately affected and severely affected *lmod1a*/*lmod1b* co-injected fish. Moderate to severe tail curvature (yellow arrows), pericardial edema (blue arrows), reduced eye size (green arrows) and abnormal distribution of melanocytes (white arrows), abnormal pooling of blood (red arrow) can be seen. **B-** Fluorescent microscopy of phalloidin stained Wild Type un-injected control (WT UIC) and *lmod1a*/*lmo1b* morpholino (MO) injected fish. Top left-WT UIC showing aortic arches (white arrows). Top right lmod1a/lmo1b MO injected fish showing aortic arches (white arrows). Bottom left-WT UIC fish showing normal muscle fibers in the tail.Bottom right-lmod1a/lmo1b MO injected fish showing crossing over of muscle fibers. **C-** Fluorescent microscopy of dextran injected WT UIC and *lmod1a*/*lmo1b* MO injected fish. Top left-WT UIC showing aortic arches (white arrows). Top right lmod1a/lmo1b MO injected fish showing aortic arches (white arrows). Bottom left-WT UIC fish showing normal vasculature in the tail. Bottom right-*lmod1a/lmo1b* MO injected fish showing absent flow towards the caudal half of the trunk, including fin vasculature. **D-** Confocal microscopy imaging of un-injected kdrl: GFP zebrafish compared with *lmod1a/lmod1b* MO injected kdrl: GFP fish. Normal branching of the pharyngeal arches can be seen in the un-injected wild-type fish but were disturbed/missing in the *lmod1a/lmod1b* MO injected kdrl: GFP fish (white arrows). Scale bar shown in the bottom right of images (100um length). *-Marks the eye of the zebrafish.

Initially, injection of one splice-directed *lmod1a* MO at a 4ng concentration generated a 67% combined moderate/severe affected rate and this increased to 93% when only *lmod1b* MO was injected at the same concentration. Two splice-directed MO (*lmod1a* and *lmod1b*) were co-injected at 4ng (2ng *lmod1a* and 2ng *lmod1b*) as a final concentration giving higher percentage of affected embryos (97%), as well as a higher survival rate (figure 4). This may suggest that as both *lmod1a* and *lmod1b* are expressed in zebrafish, when one is knocked-down, the other may have a compensatory effect. Therefore, knocking-down both paralogs more closely resemble the human deficient *LMOD1*. A human beta-globin standard control was used and injected at a 4ng concentration and the resulting fish revealed no increase in abnormal phenotype associated with toxicity compared with un-injected controls. *p53* MO was injected as an apoptosis management control at 6ng concentration and the fish did not present with any abnormal phenotype. A co-injection of *lmod1a/lmod1b/p53* MO at a final concentration of 6ng revealed no reduction in severity of the phenotype observed compared to *lmod1a/lmo1b* only co-injection. Therefore, the data presented suggests that the phenotype observed was unlikely due to off-target effects.

**Figure 4.**
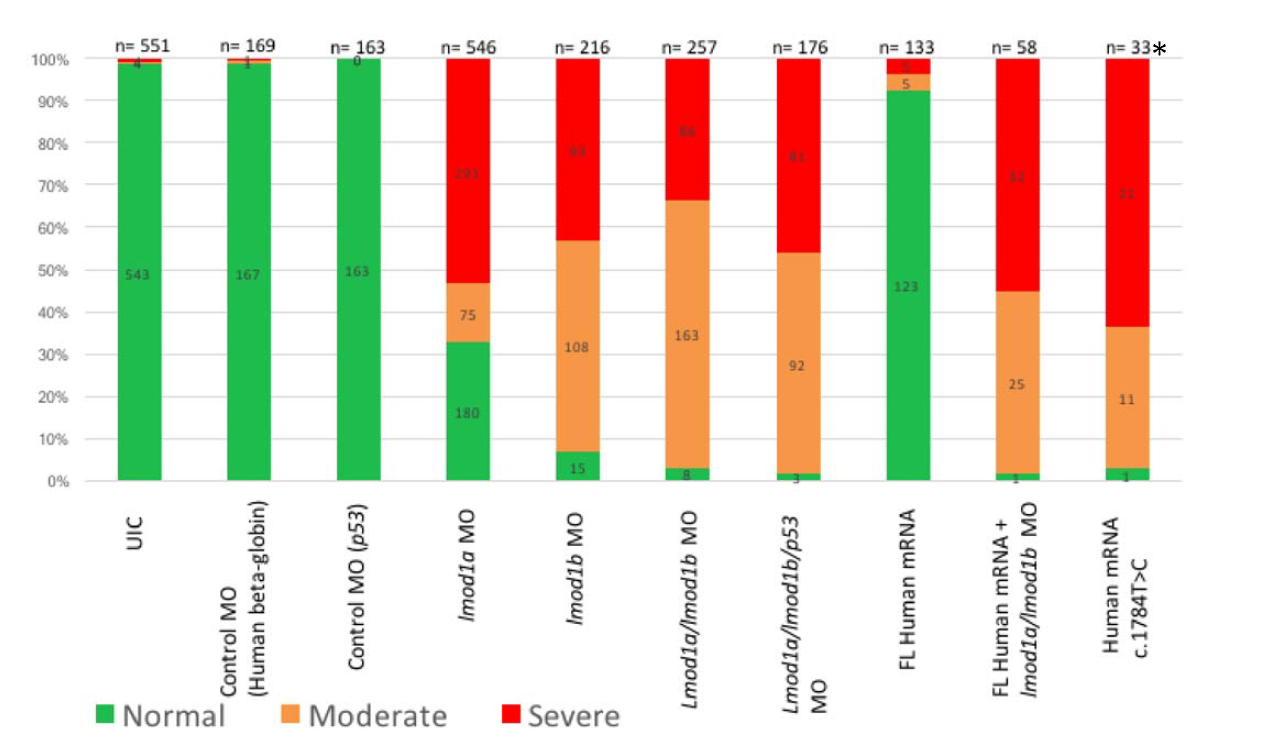
Bar chart showing normal (green), moderate (yellow) and severely affected (red) embryos associated with the following injections; Un-injected control (UIC); Human beta-globulin control MO (4ng/ul); p53 toxicity control MO (6ng/ul); *lmod1a* MO (4ng/ul); *lmod1b* MO at (4ng/ul); *lmod1a/lmod1b* MOs combined (4ng/ul; 2ng/ul + 2ng/ul); lmod1a/lmod1b/p53 MOs combined (6ng/ul; 2ng/ul + 2ng/ul + 2ng/ul); Full length *LMOD1* Wild-type human mRNA (60ng/ul); Full length *LMOD1* Wild-type human mRNA (60ng/ul) and *lmod1a/lmod1b* (4ng/ul, 2ng/ul +2ng/ul); Human mutated *LMOD1* mRNA (c.1784T>C), * A significant number of human mRNA c.1784T>C injected fish died by 24hpf compared to UIC.

Fifty out of sixty-six *lmod1a/lmod1b* MO injected embryos analysed (76%) demonstrated pooling of blood in the yolk sac by 4 days post fertilization (dpf) and as a consequence developed significant oedema in their yolk sac by 5dpf.

Injection of the full-length human wild-type *LMOD1* mRNA at 60ng/ul into 1-cell embryos did not give an overexpression phenotype and demonstrated a similar effect (7% affected) as in the standard control MO and un-injected control experiments indication no toxic effects at that concentration were causing a phenotype (figure 4).

Only the full-length mRNA of the human c.1784T>C mutation (60ng/ul concentration) was injected into the 1-cell embryos, which resulted in embryonic death in 141/174 (81%) of cases, with the remaining 32/33 zebrafish demonstrating oedema, tail curvature, abnormal distribution of melanocytes, smaller eyes and body size as seen in the MO injected moderate to severe phenotype.

Next a full length wild-type human *LMOD1* mRNA was injected at 1-cell stage followed by *lmod1a*/*lmod1b* MO (4ng/ul) and the resulting phenotype from MO knock down could not be rescued at 60ng/ul dosage or even with an increased dosage up to 200ng/ul, as most were still affected (figure 4). Wild-type zebrafish *lmod1a* and *lmod1b* were also injected singularly and combined with MO but also did not show significant rescue of the phenotype. This suggested that *LMOD1* mRNA may be required at specific time point which is not possible with zebrafish injections because these are injected at the 1-cell stage.

An F-actin (Phalloidin) marker was used to determine if knockdown of *lmod1a/lmod1b* in zebrafish had any effect on the actin filament production in the vasculature as well as the aortic arches at 4dpf (figure 3B). Crossing-over of muscle fibres through adjacent somites was also identified and disturbance/loss of the ventral pharyngeal arches was observed in the morphant fish compared to controls suggesting that *lmod1a* may be required for aortic arch development or F-actin stabilisation.

Next a high molecular weight (500kDa) fluorescein-conjugated Dextran dye was used to highlight the vasculature of the *lmod1a*/*lmod1b* injected morphants to assess for perturbations as observed in the study family. Fluorescent microscopy showed abnormal branching of the aortic arches and intermittent flow through the vessels and absent flow towards the caudal half of the trunk, including fin vasculature (figure 3C). This suggested that *lmod1a* might play a role in vascular development and maintenance.

Transgenic fish were used to further describe the abnormalities observed in the aortic arches by injection of *lmod1a*/*lmod1b* MO into *Tg(kdrl:GFP)* fish and found that MO injected fish showed minimal branching of the aortic arches compared to that of the un-injected control (figure 3D). These arches are evolutionary important because in fish, they later become the vasculature of the gills, but in humans they elaborate more extensively to carry blood to the lungs and form the carotid arteries, subclavian arteries, aorta and other associated structures in the upper mediastinum and proximal upper limb and neck. Importantly, the resulting phenotype seen in all affected fish from *lmod1a* and *lmod1b* MO as well as the mRNA of the human c.1784T>C mutation injections were seen consistently through repeat injections and toxicity controls indicate that the phenotype recorded was the result of a non-functioning protein rather than off-target toxic effects.

### Discussion

Leiomodin-1 (LMOD1) is a 64-kDa actin-binding protein and a homolog of the only known tropomyosin-coated pointed-end capping tropomodulin protein (TMOD) family^55^. Vertebrate Lmods consist of 3 isoforms; *LMOD1* (smooth muscle), *LMOD2* (cardiac muscle) and *LMOD3* (skeletal muscle). LMOD1, is expressed predominantly in human arteries (tibial, aorta and coronary), oesophagus, colon, bladder and uterus (GTEx Analysis Release V6 (dbGaP Accession phs000424.v6.p1)^56^.

No mutations in *LMOD2* have been reported yet but the Lmod2 knockout mouse exhibits shortened cardiac actin filaments and dilated cardiomyopathy^57^. Mutations in the human *LMOD3* are reported to cause thin filament disorganization and nemaline myopathy^58^ associated with the rod-like nemaline bodies reported to cause muscle weakness. The Lmod3-null mouse showed skeletal muscle weakness associated with nemaline bodies^59; 60^.

An autosomal recessive mutation causing a premature terminating codon in exon 2 of *LMOD1* was recently described as causing a fatal rare condition, megacystis microcolon intestinal hypoperistalsis syndrome (MMIHS) in a newborn child^61^. Knock-down of human LMOD1 in intestinal smooth muscle cells showed decrease in actin filament formation and contractility.

Phalloidin staining showed consistently a lower concentration of actin filaments in the Lmod1-null bladder tissue and electron microscopy revealed a presence of rod-like structures in the smooth muscle, similarly found in the Lmod3-null mouse. The authors also reported attenuation in the transitional epithelium of the bladder and stomach in the Lmod1-null mouse but not in the intestine and suggested that it may be due to decreased mechanical stress.

The data demonstrated that *LMOD1* has significant importance in regulating smooth muscle cytoskeletal-contractile coupling. In contrast to TMODs, leiomodin proteins also harbor a C-terminal extension and in all three isoforms this consists of a proline-rich region followed by a WH2 domain^62^. Recent studies have shown that constructs of *Lmod1* lacking this C-terminal, containing proline-rich region and WH2 domain, caused a delay in actin polymerization and increased dependence of *LMOD1* concentration relative to the polymerization rate^63^.

Many of the proteins known to regulate actin dynamics contain a WH2 domain, such as the β-thymosin^64^ family of proteins, and WASP proteins bind the actin nucleating protein complex Arp2/3^65^.

These data above suggest that the mutations found in the C-terminal extension containing the actin-binding domain (WH2) of LMOD1 may delay actin polymerization and therefore compromise the length and dynamics of actin filaments.

The non-synonymous *LMOD1* variant [c.1784T>C/p.(V595A)] found in family M441 changes the amino acid at the WH2 domain at residues 575-600^63^ of the protein. The mutant residue is smaller than the wild-type residue, which may lead to loss of interactions with other molecules^66^. Changes in the structure of the WH2 domain may be interfering with binding and assembly of actin filaments and therefore disrupting the actin dynamics and interaction with myosin in the smooth muscle cell contraction pathway. Delayed actin polymerization by abnormal *LMOD1* seen in other studies^63^ may lead to earlier capping of the pointed actin filaments by tropomodulin, resulting in shorter actin filaments.

The *LMOD1* c.1631C>A, p.(P544H) allele found in the proline-rich region is predicted to bind profilin, which is thought to be involved in actin dynamics, namely actin polymerization^67^. The mutant residue is larger than the wild-type, which may lead to disturbances in multimeric interactions^66^ and the hydrophobic interactions in the core or surface of the protein may become lost. Protein structure stability prediction tools forecast a decreased stability of the protein structure in both c.1631C>A [p.(P544H)] and c.1784T>C [p.(V595A)] alleles^68^.

Therefore as proline-rich regions have been reported to be contributing to actin binding alongside the WH2 domain^69^, mutations in this polyproline domain may account for a TAAD phenotype as seen in the second proband, individual M508.1.

The incidence of *LMOD1* disease-related mutations in our UK study is ˜2% (2/99) and the phenotypic presentation of this gene may be of late onset, as observed by the mean age of dissection in males (50.4 years, range 39-58 years) and variable expression of this dominant gene between carrier males and females in our study family. Of 18 adults over the age of 18 (7M:11F) carrying the c.1784T>C allele, those six with demonstrated aneurysm were predominantly male (5M:1F). Three further variants in our international cohorts (two in USA cohort and one in French cohort) were classed as likely pathogenic and further segregation studies and animal modelling are needed to confirm these.

Injection of the human mutant mRNA from the proband of the study family into zebrafish embryos may demonstrate either a dominant negative effect or haploinsufficient state. The human mutation illustrates a ’loss of function’ as the resulting phenotype is similar to that observed in *lmod1a*/*lmod1b* ‘knock down’ in zebrafish.

The likely dominant negative phenotype suggested by the animal studies indicates a loss of function mutation carried by our study family. The mutation is likely to cause a delay in the nucleation of actin monomers to actin filaments because of a non-functioning WH2 domain, resulting in decreased actin filaments^61^. Therefore, the crosslinking assembly of actin filaments may be impaired as abnormal distributions and lower concentrations of phalloidin staining have been observed by us and *Halim et al*^61^, which may consequently be leading to aortic cystic medial necrosis and permitting dilatation and dissection in the affected individuals of our study family.

The abnormal branching of the aortic arches observed in the *Tg(kdrl:GFP)* zebrafish may become an aneurysm specific gene phenotype as it has now been reported in a third TAAD gene following the *MAT2A^20^* and *FOXE3^34^ reports*.

## Conclusion

In summary, we have shown that mutations in the C-terminal end of *LMOD1* may underlie TAAD with an incidence of ˜2% in our cohort. Additional mammalian *in-vivo* knock-down studies may help to further confirm a role of LMOD1 in actin dynamics of the aorta. Histopathology will be required to explore the pathological effects (such as demonstration of cystic medial necrosis) of these mutations on the vascular smooth muscle wall in TAAD. An associated phenotype may be established, when more families and individuals with pathogenic alleles have been found. This could include blue sclerae and small joint hypermobility, features that were seen significantly more frequently in the c.1784T>C carriers than in the 98 other TAAD probands.

*LMOD1* should be investigated in TAAD exome profiles already available internationally and it may be useful to add this small three-exon gene to future TAAD panels.

Sequencing of other TAAD cohorts will determine a worldwide incidence. In addition, screening other genes in this important actin-myosin pathway may highlight other TAAD candidate genes. This should lead to more specific collaborative therapeutic trials once enough affected *LMOD1* deficient patients have been identified internationally.

## Acknowledgements

The authors would like to thank the families participating in this research study. This work was supported with grants from the British Heart Foundation to E.R.B, The Rosetrees Trust to Y.B.A.W. and A.C, The Marfan Trust, The Marfan association and The Everett Roseborough Legacy to Y.B.A.W, Peter and Sonia Field Charitable Trust to A.C, St George’s Medical School and NHS Hospital Trust to A.C. and Guy’s and St Thomas’s charity to A.S. The zebrafish study was conducted in collaboration with Zebsolutions.

